# Connecting Covert Attention and Visual Perception to the Spatiotemporal Dynamics of Alpha Band Activity, Cross-Frequency Coupling (CFC), and Functional Connectivity using Multivariate Pattern Analysis (MVPA)

**DOI:** 10.1101/2022.05.27.493754

**Authors:** Sarah S. Sheldon, Alona Fyshe, Kyle E. Mathewson

## Abstract

Some evidence suggests that alpha activity is directly related to the baseline firing rate of sensory neurons which alters the probability of responding to a stimulus. Other evidence points to alpha indirectly modulating cortical excitability through its interactions with gamma oscillations. A third possibility is that alpha-based functional connectivity better explains attentional modulation and perceptual responses. To test this, alpha amplitude, CFC, and functional connectivity measures were extracted from EEG data recorded while participants performed a cued orientation perception task. Using pre-target data, the spatiotemporal activity of each metric was submitted to a SVM classifier to determine which activity pattern best distinguished trials with covert attention from trials without. The same metrics were submitted to SVR to find the activity that best predicted task performance. Results indicate the best metric for classifying trials with and without covert attention was alpha amplitude. This indicates that, prior to target onset, alpha amplitude alone is most sensitive to the presence of covert attention. In contrast, none of the metrics were strong predictors of task performance. Overall, our results support the idea that alpha activity is directly related to changes in the baseline firing rate of sensory neurons which changes responsiveness but not performance.

---

Electroencephalography (EEG) is one of the oldest and most well-established neuroimaging techniques used by researchers and clinicians. Alpha oscillations (8-14 Hz) were the first EEG frequency band identified (Berger 1929) and are particularly interesting to cognitive neuroscience researchers due to a plethora of evidence suggesting it has a critical role in visual perception and attention (Klimesch et al. 2007; Mathewson et al. 2011; Klimesch 2012; Mathewson et al. 2012; Clayton et al. 2015; Frey et al. 2015; VanRullen 2016). Exactly what that role is, however, is still debated. For example, some researchers believe alpha oscillations inhibit the flow of irrelevant information into sensory areas or from sensory areas to higher order cortical locations (Klimesch *et al*. 2007; Jensen and Mazaheri 2010; Foxe and Snyder 2011; Mathewson *et al*. 2011; Chaumon and Busch 2014), while others think alpha oscillations indirectly control perception and attention by modulating the higher frequencies (*e.g.,* gamma oscillations, >30 Hz) involved in sensory processing (Voytek 2010; Jensen et al. 2014; Zazio et al. 2020). The purpose of the current research is to try elucidating the role of alpha oscillations in visual perception and attention using the powerful new techniques offered by machine learning. It is now thought that one of the reasons the relationship between alpha oscillations and perception remains elusive after almost a century of research is that alpha’s role in conscious perception has often been mistaken for sensory processing. For example, recent studies investigating the effect of prestimulus power on perception found that participants were more likely to report seeing a stimulus when prestimulus alpha (and low beta) power was low, even though the stimulus was not presented (Samaha et al. 2017; Iemi and Busch 2018; Samaha et al. 2020). To account for these findings, Samaha et al (2020) proposed the baseline sensory excitability model (BSEM) which states that spontaneous alpha amplitude indiscriminately changes the baseline firing rate of sensory neurons such that the sensory response distribution is changed without changing the separability of the signal and noise representations. Furthermore, because it is thought that the internal fluctuations in sensory neuron excitability are not known by the higher-level decision-making areas (Samaha *et al*. 2020), participants will fail to adjust their decision criteria accordingly so that participants are not only more likely to report stimulus presence and make more false alarms, but will also claim greater level of confidence in their decision even though their detection and discrimination sensitivity does not change. These predictions were confirmed by two studies designed for signal detection theory analysis (Samaha *et al*. 2017; Iemi and Busch 2018). Further support comes from Benwell et al (2018) who found that pre-stimulus alpha power correlated with participants’ bias when judging the relative length of two line segments (landmark task) but not discrimination performance. Interestingly, the bias seemed to be partially driven by changes in pre-stimulus alpha power over the course of the experiment rather than trial-by-trial variability.

In contrast, another group has put forth the oscillation-based probability of response (OPR) model which states that the effects of alpha oscillations on perception are due to an alpha-gamma interaction modulating the probability of sensory neurons responding to a stimulus (Zazio et al. 2020). Specially, the oscillation-based probability of response (OPR) model proposes that there are two possible alpha-gamma cross-frequency interactions that effect the probability of responding to a visual stimulus: (1) alpha power modulates the power of gamma (amplitude–amplitude coupling; suppression of gamma amplitude), so that increased alpha power results in an increased threshold for responding (Chaumon and Busch 2014); or, (2) alpha activity modulates the phase of gamma (phase–amplitude coupling,; desynchronization of gamma), so that increased alpha activity results in a decreased sensory response (Chaumon and Busch 2014). In the current study, if amplitude–amplitude coupling (AAC) activity is responsible for attentional modulation and visual perception, we would expect to see results like those proposed by the baseline sensory excitability model (BSEM) with regards to attentional effects, but a difference in how well alpha activity relates to task performance since the baseline sensory excitability model (BSEM) hypothesizes that alpha fluctuations do not change discrimination sensitivity. On the other hand, we would expect phase–amplitude coupling (PAC) to be strongly related to task performance and sensitive to the presence of an informative cue since it would modulate the signal to a larger extent than the noise. These ideas are supported by independent lines of research into the role of gamma oscillations for sensory enhancement (Pritchett et al. 2015; Ni et al. 2016) and the inverse relationship between alpha activity, cortical excitability, and sensory perception (Samuel et al. 2018; de Graaf et al. 2020), as well as research using transcranial stimulation that found stimulation at alpha frequency modulated gamma oscillation activity (Hamidi 2009; Herring et al. 2019; Okazaki et al. 2020). A summary of the hypotheses based on the two models can be found in Table 1.

**Table 1.**
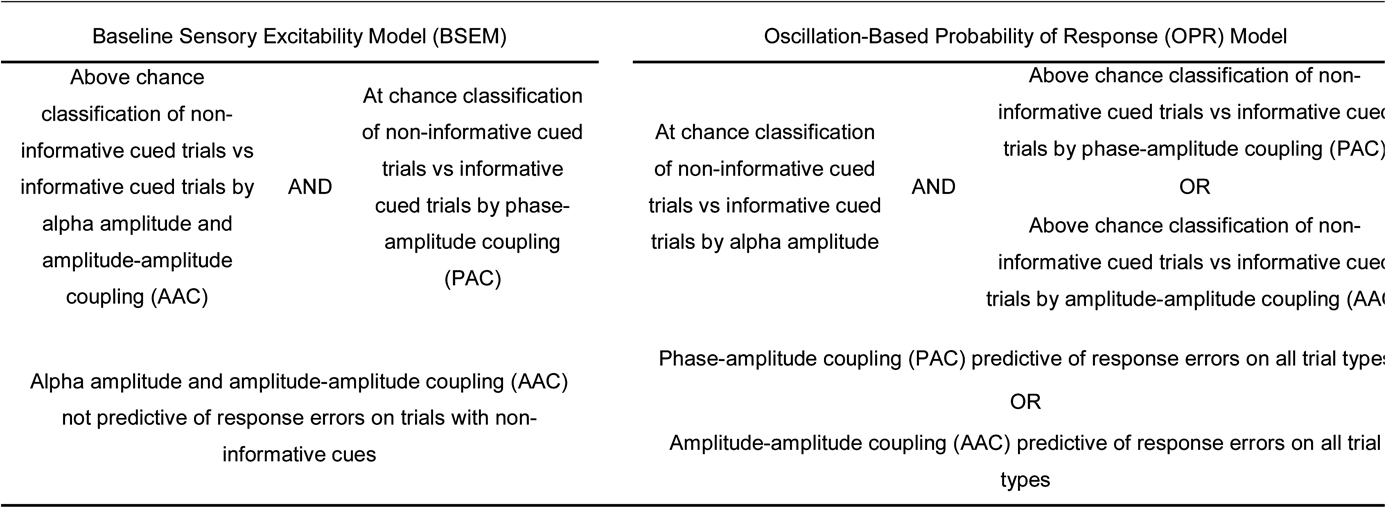
Overview of predicted outcomes that support the baseline sensory excitability model (BSEM) or support the oscillation-based probability of response (OPR) model.

In addition, rather than localized changes in brain activity, there is the possibility that coordinated activity across different brain areas might better explain attentional modulation and perceptual responses. Previous studies have reported alpha-mediated functional connectivity with regards to visual task performance and top-down attention. For example, authors found stimulus anticipation was related to occipital and medial frontal interactions in the alpha band and that participants with stronger network interactions were more likely to improve performance following an error (Cohen and van Gaal 2013). Other studies have found alpha-mediated interactions between the parietal, frontal, and occipital areas during the control of task-related attentional processes (Plomp et al. 2015; Doesburg et al. 2016). These results suggests that alpha band activity might mediate top-down modulation of visual areas not only locally, but through the dynamic interactions of large-scale functional networks (Sadaghiani and Kleinschmidt 2016).

It is important to note that the baseline sensory excitability model, the oscillation-based probability of response model, and functional connectivity are not necessarily mutually exclusive. It is possible that the oscillation-based probability of response model is a mechanistic explanation of the baseline sensory excitability model which is a result of coordination across different brain areas. It is also important to point out that neither model claims to explain attention, though both propose that their models could extend to, or are at least related to, attentional mechanisms. This leaves us with some interesting questions that need to be addressed. Namely, whether alpha power alone (baseline sensory excitability model), alpha modulated gamma (oscillation-based probability of response model), or functional connectivity best account for perceptual performance. Also, this paper sought to investigate if the same pattern of brain activity that accounts for perceptual performance can predict performance when participants are actively engaged in visuospatial attention? To address these questions, the current study uses the combination of machine learning technique, EEG data collected from a combined orientation perception and spatial attention task and associated behavioral measures (see below for further details about the task). The idea is to determine whether alpha power, alpha-gamma cross-frequency coupling, or alpha-based functional connectivity could accurately classify trials as those with selective spatial attention (informative cue) and those without (non-informative cue). In addition, the current study would also test which set of EEG measures could predict perceptual performance on a cued orientation perception task.

## Materials and Methods

### Participants

Thirty-five participants from the University of Alberta community participated in the study (age range = 17-34 years). Seven participants were not included in the analysis due to more than 30% of trials containing eye movement artifacts (see the EEG Analysis section for more details). Participants were all right-handed and had normal or corrected normal vision and no history of neurological problems. All participants gave informed written consent, were either compensated at a rate of $10/hr or given research credit for their time. The study adhered to the tenets of the Declaration of Helsinki and was approved by the Internal Ethics Board at the University of Alberta.

### Cued Orientation Perception Task

Participants were seated 57 cm away from a 1920 x 1080 pixel^2^ ViewPixx/EEG LCD monitor (VPixx Technologies, Quebec, Canada) with a refresh rate of 120 Hz, simulating a CRT display with LED backlight rastering. The rastering, along with 8-bit digital TTL output triggers yoked to the onset and value of the top left pixel, allowed for submillisecond accuracy in pixel illumination times, which were confirmed with a photocell prior to the experiment. Stimuli were presented using a Windows 7 PC running MATLAB R2012b with the Psychophysics toolbox (Version 3; Brainard 1997; Pelli 1997). The code running the task was a modified version of the Orientation Perception Task code from Sheldon and Mathewson (2021). The original version of the code can be found here: https://github.com/APPLabUofA/OrientTask_paper. Video output was sent to the ViewPixx/EEG with an Asus Striker GTX760 (Fremont, CA) graphics processing unit.

Each trial began with a white fixation dot presented at the center of the monitor and two dark gray circles to the left and right of the fixation dot for 700 ms after which one of three possible cues appeared above the central fixation dot, vertically aligned to the top of the circles. Two cues were black triangles pointing towards the left or right, indicating the side the target will appear (*i.e.,* informative cues). The third cue was both black triangles pointed toward each other indicating that the target will appear to the left or right (*i.e.,* non-informative cue). The cues remained on screen for 1242, 1284, 1325, or 1367 ms. After the cue, the target appeared for 8.33 ms (one monitor refresh) in the center of the left or right circle. The target always appeared to the side indicated by the informative cue. For non-informative cues, the target could appear on the left or right with equal probability. The target was in the shape of a needle and was pointing toward one of 24 predefined evenly spaced directions so that all the orientations covered 360 degrees. The direction of the target was randomly selected on each trial. A backward mask lasting for 8.33 ms with a constant 41.7 ms target-mask stimulus-onset asynchrony appeared in the center of both circles regardless of the side the target appeared. The mask was created by overlaying the target orientated in all 24 directions which created a star shape seen in Figure 1A. Following the mask offset, a 516.6 ms blank interval period occurred identical to the fixation. After the blank interval, a response screen appeared with the needle in the center of the screen. Using the computer mouse, participants were asked to rotate the needle so that it was pointed in the same direction as the previous target. If participants detected a target but could not remember its orientation, they were asked to guess the orientation of the target. Participants could provide their response at their own pace. No feedback was given to participants. The next trial began immediately after a needle’s orientation was selected. See Figure 1A for a summary of the task sequence and the stimulus dimensions.

**Figure 1.**
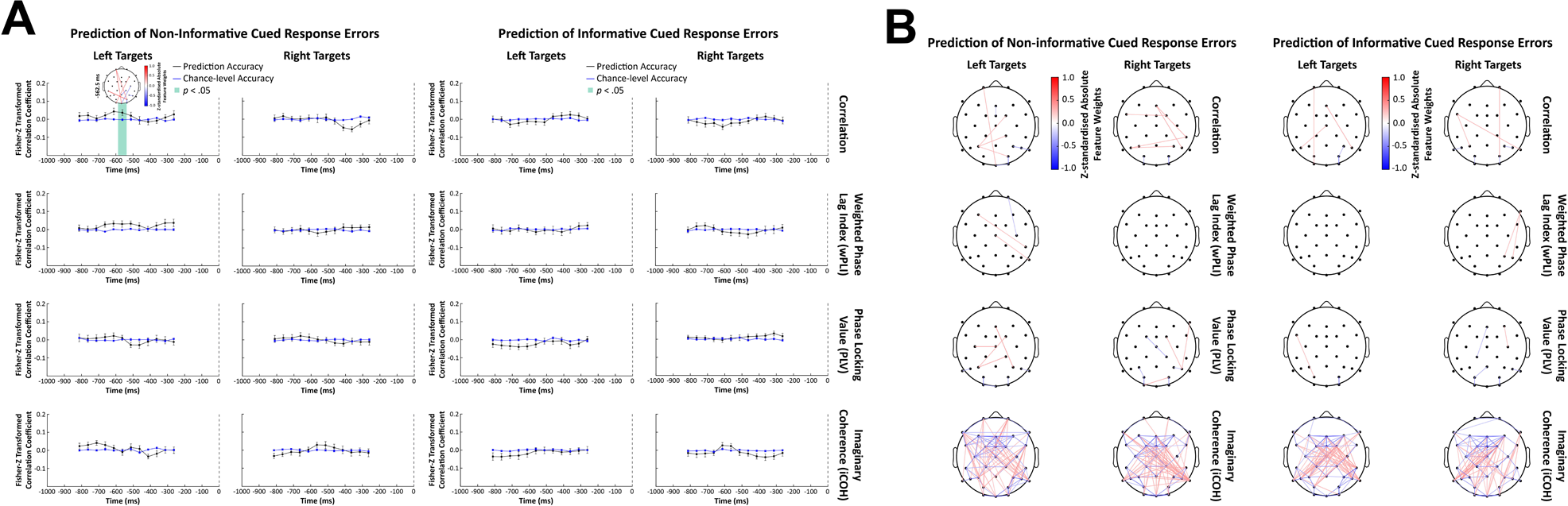
Cued orientation perception task. **A)** Sequence of task events with duration of each screen presentation and sizes of fixation, cues, target, mask, and response stimuli. Sizes are in degrees of visual angle. **B)** Example of response error calculation. Response errors are reported in degrees.

Participants completed eight blocks consisting of 48 trials each, along with 20 practice trials at the beginning of the experiment. Participants ended completing a total of 404 trials with 33% of trials having a non-informative cue and about equal left and right targets, 33% of trials having a right target and informative cue, and 33% of trials having a left target and informative cue. Participants could rest at their own pace every 48 trials. Extensive written and verbal instructions were presented to participants prior to the practice trials. Instructions thoroughly explained and demonstrated each component that would compose a single trial.

Before the cued orientation perception task, participants performed a staircased cued target detection task that had the same parameters as the cued orientation perception task except that participants only reported whether they saw the target or not using the keyboard. The target color was a gray determined by a scalar value passed to the functions in Psychtoolbox. In the staircased cued target detection task, the target color value could range from the background color (making it not visible; corresponding value of 256) to black (making it the most visible; corresponding value of 0). This target gray value was adjusted throughout the task based on a 1- up/2-down staircasing procedure targeting a 0.65 target detection rate for each individual (Garcı a-Pérez 1998; Kingdom and Prins 2016). The staircased task consisted of three blocks of 48 trials. The target gray value was determined for each participant by taking the average target gray value across the last two blocks of trials. These final values came to be 56 on average (*SD* = 20) and were used as the target gray value in the cued orientation perception task.

The MATLAB code for the cued staircased target detection task and the cued orientation perception task are available at https://github.com/APPLabUofA/CuedOrientTask_paper.

### EEG Recording

During the experiment, EEG data was recorded from each participant with a Brain-Amp 32-channel amplifier (BrainVision) using gelled low-impedance electrodes (actiCAP passive electrodes kept below 5 kΩ). Inter-electrode impedances were measured at the start of each experiment. All electrodes were arranged in the same 10-20 positions (Fp2, Fp1, F4, F3, F8, F7, FC2, FC1, FC6, FC5, C4, C3, CP2, CP1, CP6 CP5, P4, P3 P6, P5, P8, P7, PO4, PO3, O2, O1, Fz, FCz, Cz, Pz, and Oz). In addition to the 31 EEG sensors, a ground electrode was used, positioned at AFz. Two reference electrodes and the vertical and horizontal bipolar EOG were recorded from passive Ag/AgCl easycap disk electrodes affixed on the mastoids, above and below the left eye, and 1 cm lateral from the outer canthus of each eye. The bipolar channels were recorded using the AUX ports of the Brain-Amp amplifier. SuperVisc electrolyte gel and mild abrasion with a blunted syringe tip were used to lower impedances. Gel was applied and inter-electrode impedances were lowered to less than 5 kΩ for all electrode sites. EEG data was recorded online referenced to an electrode attached to the left mastoid. Offline, the data were re-referenced to the arithmetically derived average of the left and right mastoid electrodes.

Data were digitized at 1000 Hz with a resolution of 24 bits. Data were filtered with an online bandpass with cutoffs of 0.1 Hz and 250 Hz. The experiment was run in a dimly lit, sound and radio frequency-attenuated chamber from Electromedical Instruments, with copper mesh covering the window. The only electrical devices in the chamber were an amplifier, speakers, keyboard, mouse, and monitor. The monitor ran on DC power from outside the chamber, the keyboard and mouse were plugged into USB outside the chamber, and the speakers and amplifier were both powered from outside the chamber, and nothing was plugged into the internal power outlets. Any devices transmitting or receiving radio waves (e.g., cell phones) were removed from the chamber for the duration of the experiment.

### Data Analyses

Data analyses were performed using MATLAB R2021a (The MathWorks Inc, Natick, MA, USA), EEGLAB 14.1.2b (Delorme and Makeig 2004), FieldTrip 20160928 (Oostenveld et al. 2011), Python 3.7.9 (Python Software Foundation, https://www.python.org/) via the Spyder IDE (version 5.0.5; Raybaut 2009), and custom scripts. All statistical analyses were conducted using MATLAB R2021a. The functional connectivity topographic plots were made with code adapted from the FCLAB toolbox (Pezoulas et al. 2018) found here: https://github.com/ramsys28/FCLAB. The raw behavior and EEG data files are available at https://osf.io/u6fgm/ and the analysis code is available at https://github.com/APPLabUofA/MVPA_CuedOrientTask_paper.

#### Behavioral Data

Response errors on each trial were calculated by subtracting the orientation of the response stimulus, as reported by the participant, from the orientation of the target stimulus (see Figure 1B). The response errors and absolute response errors were submitted to a repeated-measures factorial ANOVA (target side and cue type the within-subject factors) using the permutation-based *Fmax* test which is a mass univariate approach for factorial ANOVAs. The *Fmax* test has the advantages of permutation tests in there are a lot less assumptions about the data than traditional statistical approaches, and has been shown to be very good at controlling Type I errors without a great cost in power (Fields and Kuperberg 2019). The implementation of the *Fmax* test was done with code modified from the Factorial Mass Univariate Toolbox (FMUT; Fields 2017). The original code can be found here: https://github.com/ericcfields/FMUT.

Follow-up pairwise comparisons of significant interactions and main effects were done with a repeated-measures, two-tailed permutation test based on the *tmax* statistic (Blair and Karniski 1993) using the mxt_perm1() function from the Mass Univariate ERP Toolbox (Groppe et al. 2011). *P*-values were then corrected for multiple comparisons using the false discovery rate (FDR) procedure described in Benjamini and Yekutieli (2001). Effects that satisfied a 5% FDR criterion were considered significant. 10,000 random within-participant permutations were used to estimate the distribution of the null hypothesis for all permutation-based tests (*i.e., Fmax* and *tmax*) and the familywise alpha (α) was set to 0.05.

#### EEG Data

##### Preprocessing

After the data had been re-referenced offline, the bandpass FIR filter from EEGLAB was applied with lower and upper cut-offs of 0.1 Hz and 50 Hz. Data was segmented into 3000 ms epochs aligned to target onset (−2400 ms pre-target onset to 1400 ms post-target onset). The average voltage in the 200 ms baseline following fixation onset was subtracted on each trial for every electrode, and trials with absolute voltage fluctuations on any channel greater than 1000 μV were discarded. Eye movements were then corrected with a regression-based procedure developed by Gratton, Coles, and Donchin (1983). After a second baseline subtraction with 200 ms following fixation onset, trials with remaining absolute voltage fluctuations on any channel greater than 500 μV were removed from further analysis.

Data was then subjected to visual inspection and manual rejection of trials (both EEG and behavioral data) contaminated by blocking (*i.e.,* amplifier saturation), blinks, muscle noise, or skin potentials. To ensure there were no horizontal eye movements during the trial, we use a split-half sliding window approach on the HEOG signal. We slid a 100 ms time window in steps of 10 ms from 1600 ms prior to target onset (at least 200 ms before the first cue appears) to the response screen onset 600 ms after target onset. If the change in voltage from the first half to the second half of the window was greater than 25 µV, it was marked as an eye movement and rejected (Hong et al. 2015; Hakim et al. 2019). Seven participants had to be removed from further analysis after this procedure because more than 30% of trials got rejected indicating that they did not use covert attention as instructed.

##### Periodic and Aperiodic Components

Because neural power spectra consist of overlapping periodic and aperiodic components, it is important to consider the aperiodic activity when measuring a signal’s oscillatory properties. For this reason, the FOOOF toolbox (version 1.0.0; Donoghue et al. 2020) was used to identify peaks within the alpha (8-14 Hz) frequency range after adjusting for the aperiodic component. FOOOF is a spectral parameterization algorithm which decomposes the power spectrum into periodic and aperiodic components via an iterative process of model fitting (see Donoghue et al., (2020) for detailed description). The FOOOF toolbox is available on https://github.com/fooof-tools/fooof.

Before FOOOF, the power spectral density (PSD) was estimated using Welch’s method by applying the pwelch function from MATLAB’s Signal Processing Toolbox to the 1000 ms before target onset for each participant, electrode, and trial separately. The PSD was calculated using a Hamming window based on a 500 ms periodogram to perform spectral decomposition at each frequency. The PSD data was then submitted to FOOOF in Python.

Settings for the FOOOF algorithm were set as: peak width limits: [1,5]; max number of peaks: 7; minimum peak height: 0.22; peak threshold: 0.20; and aperiodic mode: fixed. Power spectra were parameterized across the frequency range 3.9 to 48.8 Hz with a frequency resolution of 1.9 Hz.

The measures of the aperiodic component were offset and exponent. The measures quantifying the quality of fit were r^2^ and fit error. All four measures were submitted to a repeated-measures factorial ANOVA (target side and cue type the within-subject factors) using the permutation-based *Fmax* statistic. The implementation of the *Fmax* test was done with code modified from the Factorial Mass Univariate Toolbox (FMUT; Fields 2017). All brain electrodes were included in the analysis 10,000 random within-participant permutations were used to estimate the distribution of the null hypothesis and the familywise alpha (α) was set to 0.0125 (Bonferroni corrected alpha level α_corr_ = 0.05/4 to account for the two aperiodic component measures and the two quality of fit measures).

##### Alpha (8-14 Hz) Peak Detection

Alpha peaks were identified on a trial-by-trial basis for each electrode and participant separately using the FOOOF algorithm. The final alpha peak frequencies used in subsequent analyses were the median frequency of the trials in each trial category (*i.e.,* informative cue, right target; non-informative cue, right target; informative cue, left target; and, non-informative cue, right target). This meant that each participant, electrode, and trial category had a peak alpha frequency that could be used in later analyses.

#### EEG Signal Metrics

All EEG metrics were calculated for each trial type, electrode, and participant separately. A summary of the extracted EEG signal measures can be found in Table 2. All features depend on both band-pass filtering and applying Hilbert transform to obtain instantaneous phase, amplitude, and frequency. Because both models are concerned with the influence of alpha oscillations on stimulus perception, only brain activity before target onset was considered. It was also important to use a time window starting at least 200 ms after the last cue onset to prevent contamination from the visual evoked potential (Luck 2014). Thus, all EEG features were calculated for the 1000 ms prior to target onset (*i.e.,* −1000 to 0 ms) with the condition that the time windows over which the metrics were calculated never exceeded 0 ms.

**Table 2.**
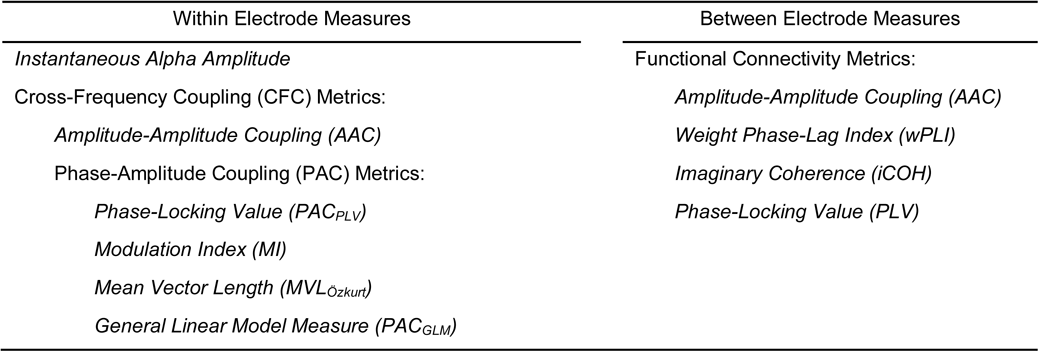
Summary of the EEG metrics used for SVM classification and prediction.

| Within Electrode Measures | Between Electrode Measures |
| --- | --- |
| *Instantaneous Alpha Amplitude* | Functional Connectivity Metrics: |
| Cross-Frequency Coupling (CFC) Metrics: | *Amplitude-Amplitude Coupling (AAC)* |
| *Amplitude-Amplitude Coupling (AAC)* | *Weight Phase-Lag Index (wPLI)* |
| Phase-Amplitude Coupling (PAC) Metrics: | *Imaginary Coherence (iCOH)* |
| *Phase-Locking Value (PAC<sub>PLV</sub>)* | *Phase-Locking Value (PLV)* |
| *Modulation Index (MI)* |  |
| *Mean Vector Length (MVL<sub>Özkurt</sub>)* |  |
| *General Linear Model Measure (PAC<sub>GLM</sub>)* |  |

To prevent edge artifacts, the data was zero-padded so that filtering was done on an additional 2000 ms (1400 ms data + 600 ms zero-padding) on either side of the selected window (Cohen 2008). The filter type used was a 4^th^ order Butterworth Infinite Impulse Response (IIR) filter implemented using Fieldtrip’s ft_preproc_bandpassfilter() function (Oostenveld *et al*. 2011). To remove any phase distortion caused by filtering in the original signal, we used a backward forward zero phase filtering method. A variable width filter was used for the cross-frequency coupling (CFC) calculations (see Cross-Frequency Coupling (CFC) section for more details), and an 8-14 Hz wide filter was used for the functional connectivity analysis. Afterward, the Hilbert transform was applied using Fieldtrip’s ft_preproc_hilbert() function (Oostenveld *et al*. 2011) or the Hilbert() function from MATLAB’s Signal Processing Toolbox. The reason both functions were utilized was that the code calculating the cross-frequency coupling (CFC) metrics originated from different sources who used either Fieldtrip’s or MATLAB’s function. In other words, when code was adapted from other sources, that function was not changed (see below for more details).

The instantaneous amplitude extracted after filtering ±1 Hz of the median peak alpha frequency was used as the metric of instantaneous alpha amplitude submitted to the decoding analysis along with the cross-frequency coupling metrics describe below. Because of the Hilbert transformation, there were 1001 time points (−1000 to 0 relative to target onset) for the alpha metric.

##### Cross-Frequency Coupling

Cross-frequency coupling metrics are used to quantify the interactions between frequency bands. There are two types relevant to the current study: phase– amplitude coupling and amplitude–amplitude coupling (Jensen and Colgin 2007; Cohen 2014; Davoudi et al. 2020). Both were calculated using a 250 ms time window, the minimum needed to capture two full cycles of the lowest frequency oscillation at the 1000 sampling rate, and a 1 ms time step.

The bandwidth of a filter used is a crucial parameter when calculating CFC. For extracting the instantaneous amplitude, filters need to be wide enough to capture the amplitude fluctuations. However, extracting the instantaneous phase requires a narrow band filter for accurate estimation (Davoudi, Ahmadi and Daliri, 2020). For this reason, a variable bandwidth, defined as ±0.4 times the center frequency was used on the faster 30-45 Hz frequency range (results in 15-60 Hz band-pass filter; Aru et al. 2015; Seymour et al. 2017), and a precise ±1 Hz of the median peak alpha frequency was used for the slower frequency range (Seymour *et al*. 2017).

Amplitude-amplitude coupling (AAC) was measured using Spearman’s rho which assess the correlation between the instantaneous amplitude of alpha and gamma (Cohen 2014). Values can range between 1 and −1 where 0 indicates no coupling and ±1 indicated complete coupling.

Since there is little agreement about the best way to quantify phase-amplitude coupling (PAC), several different metrics were chosen, each with their own benefits and limitations. For instance, the phase-locking value (which will be referred to as PAC_PLV_) described by Cohen (2008) was calculated using a modified version of the pacMEG.m function by Seymour and colleagues (2017). The PAC_PLV_ represents the degree to which the faster (gamma) oscillations are comodulated with the slower frequency’s (alpha) phase. It was originally developed as a flexible way to identify PAC across time without requiring *a priori* assumptions about the frequency bands at which coupling will be assessed (Cohen 2008). The value of 1 represents complete coupling and 0 represents no coupling.

The Modulation Index (MI), as proposed by Tort and colleagues (2008; 2010), measures cross-frequency coupling based on Shannon entropy and Kullback-Leibler (KL) divergence. When the mean amplitude is uniformly distributed over the phases (*i.e*., lack of phase-amplitude coupling), MI = 0. As the amplitude distribution deviates further from a uniform distribution, the closer MI gets to 1 indicating the existence of phase-amplitude coupling. The MI was calculated with a modified version of code from the EEGLAB extension PACTools (version 1.0.1; Martinez-Cancino et al. 2020). The original code can be found here: https://github.com/sccn/PACTools. The number of bins was 18 except when that resulted in empty bins. To avoid having empty bins (which results in a null value), the number of bins was iteratively decreased by half until none of the bins were empty.

Alternatively, the mean vector length (MVL) created by Canolty et al (2006) is calculated by multiplying slow oscillation phase time series by fast oscillation amplitude and then averaging the vectors across time. The length of the average vector represents the amount of phase-amplitude coupling and the direction represents the mean phase where amplitude is strongest. When no coupling is present, all vectors cancel each other out and the mean vector will be short (Hülsemann et al. 2019). The problem with the mean vector length (MVL) is that it can be dependent on the absolute amplitude of the high frequency oscillation (Tort *et al*. 2010). To address this caveat, Özkurt and Schnitzler (2011) proposed a direct MVL (which will be referred to as MVL_Özkurt_) which is amplitude-normalized and ranges between 0 and 1. MVL_Özkurt_ was used as a phase-amplitude coupling measure and it was calculated using a modified version of the pacMEG.m function by Seymour and colleagues (2017).

Finally, Penny et al (2008) created a measure of phase-amplitude coupling based on the general linear model (GLM). This measure, which will be referred to as PAC_GLM_, uses multiple regression to estimate (via least squares solutions) the regression coefficients of a model relating the amplitude of the faster frequency to the phase of the slower frequency (see Penny et al (2008) for details). The final PAC_GLM_ metric is the proportion of variance explained by the model (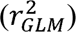) which ranges between 0 and 1. The PAC_GLM_ was meant as an improvement to the envelope-to-signal correlation measure by Bruns and Eckhorn (2004) by removing sensitivity to amplitude co-modulation and allowing coupling to be detected at all phases of the slower oscillation (Penny *et al*. 2008). The PAC_GLM_ was calculated with a modified version of code from the EEGLAB extension PACTools (version 1.0.1; Martinez-Cancino *et al*. 2020). The original code can be found here: https://github.com/sccn/PACTools.

##### Functional Connectivity

Functional connectivity metrics are used to quantify the synchronization of neural responses across electrodes. Volume conduction is a possible confound in functional connectivity analysis that can be dealt with in several ways. One method is to use measurements that are relatively insensitive to the effects of volume conduction such as phase-lag index (PLI), weighted PLI, and imaginary coherence (iCOH; Cohen 2014). These measures were calculated for all electrode pairs following the bandpass filtering and Hilbert transform.

Another method is to apply a spatial filter, like the surface Laplacian, which minimizes volume-conduction effects (Kayser and Tenke 2015). This was done prior to the bandpass filtering using the algorithm described by Perrin and colleagues (1989) and the function laplacian_perrinX.m written by Cohen (2014) which can be found at https://github.com/mikexcohen/AnalyzingNeuralTimeSeries. The function’s default settings were used which had the Legendre polynomial order was set to 20 and the G smoothing parameter (lambda) set to 1e-5. After spatial and frequency filtering and the Hilbert transform, phase-locking values (PLV) and amplitude envelope correlations were calculated for all electrode pairs. All functional connectivity measurements were calculated between −1000 ms and 0 ms (relative to target onset) in 375 ms time windows, the minimum segment length for at least three cycles of the lowest frequency band as recommended by Cohen (2014), and 5 ms time steps.

There are several other metrics used to quantify functional connectivity. There are a lot more ways to quantify functional connectivity than the measures used here, but the metrics chosen, which are summarized below, were those that best balanced computational intensity with suitability for the experimental design and research question.

Phase lag index (PLI) aims to obtain reliable estimates of phase synchronization that are unaffected by volume conduction. The central idea is to discard phase differences that center around 0 mod π since zero phase delay between two points is one of the properties of volume conduction. It is calculated as the average number of phase angle differences that are positive or negative in the complex plane (Cohen 2015). The weighted phase lag index (wPLI) is an extension of the phase lag index (PLI) proposed by Vinck *et al* (2011) where the vectors closest to the real axis are given less weight, thus, have a smaller influence on the final connectivity estimate (Cohen 2015). The wPLI in the current study was corrected for sample-size bias using the debiased estimator introduced by Vinck *et al* (2011). wPLI was calculated using modified code based on Fieldtrip’s (Oostenveld *et al*. 2011) implementation found here: https://github.com/fieldtrip/fieldtrip/tree/master/connectivity; and, online code by Cohen (2014) found here: https://github.com/mikexcohen/AnalyzingNeuralTimeSeries.

Imaginary coherence (iCOH) was developed by Nolte *et al* (2004) as a way to measure spectral coherence without the influence of volume conduction (Cohen 2014). The iCOH was calculated using a modified version of the online code provided by Cohen (2014) found here: https://github.com/mikexcohen/AnalyzingNeuralTimeSeries.

Phase-locking value (PLV) described by Lachaux *et al* (1999) measures the uniformity of the distribution of phase angle difference between two electrodes. However, instead of calculating the phase difference across trials, the phase difference between electrodes was calculated across time within each trial. So, if the phase difference varies little within a time window, the PLV is close to one. If there is a lot of variability within a time window, the PLV is close to 0. The PLV was calculated using a modified version of the pn_eegPLV.m function from the FCLAB toolbox (Pezoulas *et al*. 2018). The original code can be found here: https://github.com/ramsys28/FCLAB.

Like the cross-frequency coupling metric amplitude-amplitude coupling (AAC), the amplitude envelope correlations were calculated using Spearman’s rho because it does not rely on an assumption of normally distributed data. The only difference in the calculation was that the amplitude envelope correlations were done between electrode pairs rather than between two different frequencies at the same electrode.

### Multivariate Pattern Analysis (MVPA)

Traditionally, data analysis has primarily been limited to a univariate approach such as detecting differences in activity between experimental conditions. In contrast, MVPA is concerned with how multivariate neural patterns comprising spatial and temporal combinations might collectively correspond to a cognitive event or state of interest (Kuntzelman et al. 2021). As such, MVPA is a powerful technique to demonstrate the availability of discriminatory or predictive information, without requiring many assumptions about the underlying spatial or temporal extent of that information (Hogendoorn 2015). Another important advantage of this approach is that it does not require averaging over space, time, trials, or participants. Furthermore, MVPA can be applied at individual time points, allowing for the investigation of how the information content changes over time (Hogendoorn 2015).

A linear support vector machine (SVM) was chosen for the current MVPA. The SVM is a generalization of the maximal margin classifier developed in the 1990s by computer scientist (Vapnik 1996). Because the goal of the current study was not high prediction accuracy, the linear SVM was preferrable to the more sophisticated nonlinear algorithms because the linear SVM is simpler, making interpretation less complex (Grootswagers et al. 2017).

It should be noted that finding discriminatory or predictive information in EEG activity patterns using MVPA does not mean that those activity patterns are how the brain represents that information. In other words, it should not be assumed MVPA represents a biologically plausible mechanism employed by the brain. At best, MVPA shows that information about the experimental conditions is latent in brain activity patterns, not that the information is being used by the brain (Ritchie et al. 2019).

With regards to the current analysis, each metric was min-max normalized across trials for each participant before SVM classification and regression because it can speed up calculations and help increase accuracy (Ben-Hur and Weston 2010).

#### Support Vector Machine (SVM) Classification

The goal of the SVM classification analysis was to determine the patterns of brain activity before target onset that best distinguish between informative and non-informative cued trials (left and right targets separately). Support vector machine (SVM) classification works by finding the hyperplane which maximizes the margin between categories.

To perform classification, we used C-SVM (implemented by the LIBSVM Toolbox; Chang and Lin 2011) with a linear kernel and the regularizing/cost parameter CLJ=LJ1. This was implemented with the Decision Decoding Toolbox (DDTBOX; Bode et al. 2019) and custom script. The number of trials were balanced between the two conditions before classification because a trained classifier could achieve high accuracy in an unbalanced dataset by predicting the more frequent trial category (Grootswagers *et al*. 2017).

### Support Vector Regression (SVR)

When the variables of interest are continuous rather than categorical, an alternative to support vector machine (SVM) classification is support vector regression (SVR). SVR allows for trial-by-trial values of a continuous variable (in this case, response errors) to be mapped to predicted values of that variable (Bode *et al*. 2019). The goal of this analysis was to determine the patterns of brain activity before target onset that best predict response errors (done separately for each individual and the following four conditions: informative cue, right target; non-informative cue, right target; informative cue, left target; and, non-informative cue, left target).

Support vector regression (SVR) works differently than support vector machine (SVM) classification in that the support vectors of SVR are the data points (*i.e.,* EEG signal metrics calculated at each electrode and timepoint) with relatively large residuals (*i.e.,* furthest from the regression line) rather than the closest (Bishop 2006). However, in both cases, the data points that are hardest to categorize (closest together) or predict (furthest from their predicted values) are the support vectors used to determine the solution (Bode *et al*. 2019).

Support vector regression (SVR) analyses was implemented with the Decision Decoding Toolbox (DDTBOX; Bode *et al*. 2019) code and custom script. The SVR model was calculated using LIBSVM with a standard cost parameter C = 0.1 (Chang and Lin 2011).

### Cross-Validation

A 10-fold cross-validation procedure repeated 10 times was used to assess performance of the support vector machine (SVM) classifier/model for each participant and set of conditions. In this method, the data are divided into 10 subsets. The classifier/model is trained on nine subsets and tested using the left-out subset. This procedure is then independently repeated with each subset serving at the test data once while training on the remaining nine data sets. The average 10 cross-validation steps and 10 repetitions of the entire cross-validated analysis results in 100 analyses which was averaged, giving the final classification/prediction accuracy (Grootswagers *et al*. 2017; Bode *et al*. 2019).

For support vector regression (SVR), the only difference to the classification analysis was that the results of SVR are individual correlation coefficients between the predicted variable based on the regression model and their true values, averaged across all 100 analyses (10-fold cross-validation and 10 repetitions). Finally, the Fisher-Z transformation was applied to the correlation coefficients so that the final values could approach a normal distribution (Bode *et al*. 2019; Bode et al. 2021).

Permuted-labels analyses can be used to estimate a distribution for the null hypothesis (in this case the null hypothesis is that the data patterns associated with each category are exchangeable). This was done by repeating all original analyses with the same data and category labels, but with assignment of labels independently randomized for each iteration. The only difference between the actual classification analysis and the permuted-labels/chance analysis was that 20 repetitions were used in the permuted-labels calculations rather than 10 to generate a better approximation of the null distribution. The final statistical chance level was obtained by averaging the estimates from 200 analyses (*e.g*., 10-fold cross-validation with 20 full repetitions). Then this chance accuracy was compared with the actual classification accuracy to check the statistical significance (Bode *et al*. 2019; Bode *et al*. 2021). The same permuted-labels analyses were applied to the support vector regression (SVR) analyses for each participant and each condition to obtain a distribution of regression results under the null hypothesis.

For SVM classification and SVR, a moving time window was used to perform the cross-validation calculations and analyses over the pre-target time-period. The choice in window width and step size was based on balancing data size and temporal resolution. All the cross-frequency coupling (CFC) measures used a 15 ms time window and 15 ms step size which meant that there were 50 timepoints at each electrode. The alpha amplitude metric used a 20 ms time window and 20 ms step size which resulted 50 timepoints at each electrode. All the functional connectivity metrics used a 50 ms (10 timepoints multiplied by the step size of the original calculations) time window and 50 ms (10 timepoints multiplied by the step size of the original calculations) step size which meant that there were 12 timepoints at each electrode. It should be noted that the timepoints in the cross-validation analysis are cross-frequency coupling (CFC) and functional connectivity measurements calculated from larger time windows so there is less temporal precision than it might appear.

For group level statistical analysis, t-tests using a threshold of *p* < 0.05 were used to compare the empirical results with permuted-labels results (Bode *et al*. 2019). Subsequently, right-tailed *tmax* permutation testing (number of iterations = 10000, α level = 0.05) was conducted for multiple comparisons correction (Blair and Karniski 1993).

### Feature Weight Analysis

Support vector machine (SVM) feature weights are measures of the relative importance of each feature for classification or regression. Feature weights describe the contribution of each feature in determining the decision boundary or regression line (Bode *et al*. 2019). Importantly, a weight parameter does not reflect the contribution of each feature in isolation. Instead, the weight parameter directly reflects the usefulness of that feature to the discrimination or prediction process in the context of the other features (Hebart and Baker 2018).

For the feature weight analysis, the absolute feature weights were extracted for each electrode at time windows showing significance at the group level. Each electrode weight was calculated as the average feature weight across the time window at that channel. Raw feature weights were transformed using the method introduced by Haufe *et al* (2014) to ensure accurate topographies. Then the transformed feature weights were converted into *z*-scores and submitted to a right-tailed *tmax* permutation testing (number of iterations = 10000, α level = 0.05) for multiple comparisons correction (Blair and Karniski 1993). It is important to note that the reliability of activity patterns depends on the quality of the weights which, in turn, rely on classification/prediction performance. If accuracy is low, weights are likely suboptimal, and reconstructed activation patterns have to be interpreted with caution (Haufe *et al*. 2014; Grootswagers *et al*. 2017).

## Results

### Behavioral Data

As can be seen in Figure 2, The response errors showed no significant main effects (target side: *Fmax*(1,27) = 2.04 (critical *Fmax* = ±4.03), *p* = 0.15; cue type: *Fmax*(1,27) < 1) or interactions (*Fmax*(1,27) = 1.36 (critical *Fmax* = ±4.16), *p* = 0.25). In contrast, the absolute response errors, which can be thought of as the magnitude of the response error, had a significant main effect for target side (*Fmax*(1,27) = 13.78 (critical *Fmax* = ±4.16), *p* < 0.001), cue type (*Fmax*(1,27) = 9.80 (critical *Fmax* = ±4.27), *p* < 0.01), and a significant interaction between target side and cue type (*Fmax*(1,27) = 17.19 (critical *Fmax* = ±4.06), *p* < 0.001).

**Figure 2.**
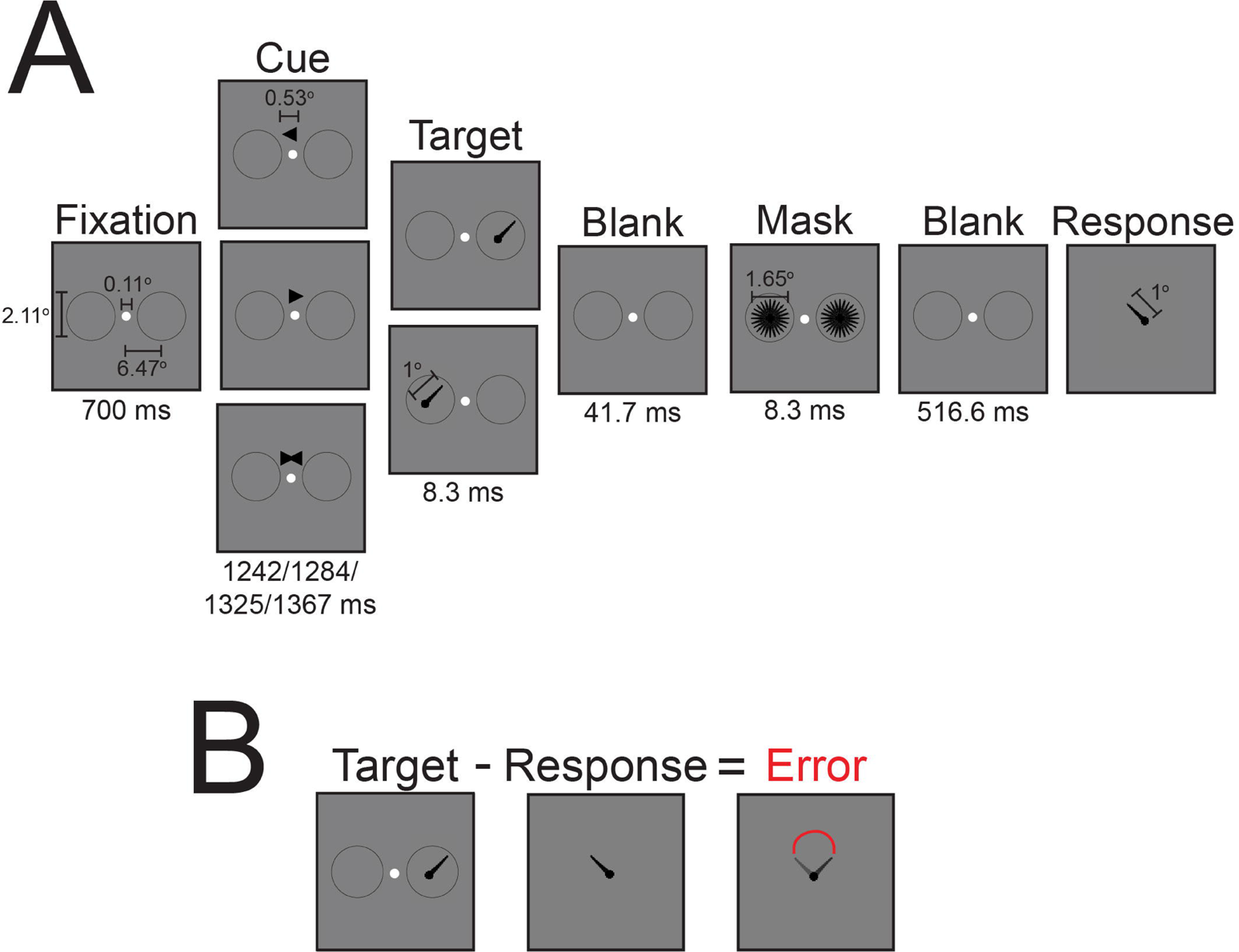
Summary of behavioral results. *Left,* significant effects of experimental conditions on the absolute values or magnitude of the response errors. Error bars are the ±*SEM. Right,* the interaction and main effects were not significant for participants’ response errors. Error bars are the ±*SEM*.

### Aperiodic Component

The aperiodic offset and exponent showed no significant main effects or interactions (mean critical *Fmax*(1,27) = +/-14.67 and mean critical *Fmax*(1,27) = +/-15.05, respectively). The quality of fit measures r^2^ and fit error also showed no significant main effects or interactions (mean critical *Fmax*(1,27) = +/-13.74 and mean critical *Fmax*(1,27) = +/-16.27, respectively).

### Support Vector Machine (SVM) Classification

As shown in Figure 3A, the SVM was able to accurately distinguish between trials that had an informative cue from those with a non-informative cue by the spatiotemporal pattern of alpha amplitude before target onset. The classification accuracy was significantly above chance at most of the time windows regardless of whether the target was to appear on the left or right. Feature weight analysis averaged over the entire epoch revealed that the major contributors to the discrimination process were the central and medial electrodes in the centroparietal, parietal, parietooccipital and occipital brain areas. However, when the targets would appear on the left, the left and central occipital electrodes were not significant contributors to the classification analysis whereas they were for when the upcoming targets would to appear on the right (Figure 3B).

**Figure 3.**
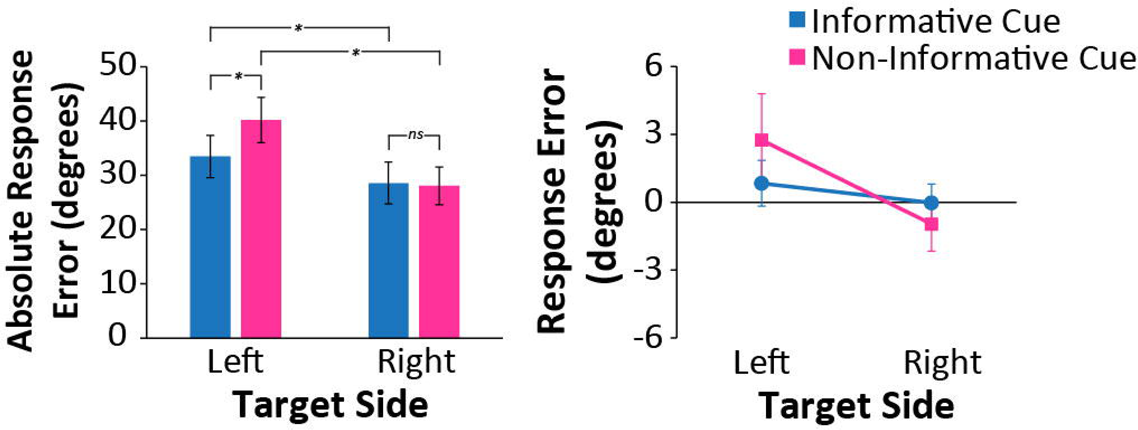
Results from spatiotemporal SVM classification and corresponding feature weight analysis based on alpha amplitude and cross-frequency coupling (CFC) measures. **A)** Spatiotemporal classification accuracy of trials with non-informative vs informative cues across all electrodes from alpha amplitude and cross-frequency coupling (CFC) measures. The SVM was trained separately on trials with left targets and trials with right targets. The black and blue lines show the actual accuracy, and the permutation test results, respectively. Error bars indicate standard error of the mean. The green shaded regions indicate the classification accuracy was significant at these time points (*p* < 0.05, corrected for multiple comparison). The dashed gray line indicates target onset. Topographic plots are the z-standardized absolute feature weights at or across the significant time points indicated by the text to the left of the heads. Green stars on the topographic plots are electrodes found to be significant features after correction for multiple comparisons. **B)** Topographic maps of the *z*-standardized absolute feature weights averaged over the −1000 ms epoch prior to target onset. Green stars indicate the electrodes with *z*-score feature weights significantly above zero after correction for multiple comparisons. Only measures with classification accuracy scores significantly above chance have stars indicating the electrodes with significant feature weight *z*-scores. AAC = amplitude-amplitude coupling; MI = modulation index; MVL = mean vector length; PAC = phase-amplitude coupling.

#### Cross-Frequency Coupling (CFC)

The only CFC measure with an activity pattern that trained the support vector machine (SVM) to classify trials significantly better than chance was the MVL_Özkurt_ (Figure 3A). For left targets, this was at −875 ms relative to target onset; for right targets, this was for the time window −785 to −770 ms and at −380 ms relative to target onset. The feature weights calculated for the entire epoch and at the significant time windows were not significant contributors to determining the decision boundary (Figure 3B). Note that because feature weights are only as useful as the information represented by the pattern of activity they are derived from (*e.g.,* the relative importance of electrode activity for classification is meaningless if there is no way to accurately classify trials in the first place), only measures that had classification accuracies above chance are discussed.

#### Functional Connectivity

The only functional connectivity measure that had classification accuracies above chance was the weighted phase lag index (wPLI). As can be seen in Figure 4A, the significantly above chance classification accuracy (black solid line) was only for targets on the right and at −462.5 ms relative to target onset. Feature weight analysis revealed that none of the electrode connections were significant contributors to the decision boundary. The same was for the feature analysis over the entire epoch (Figure 4B).

**Figure 4.**
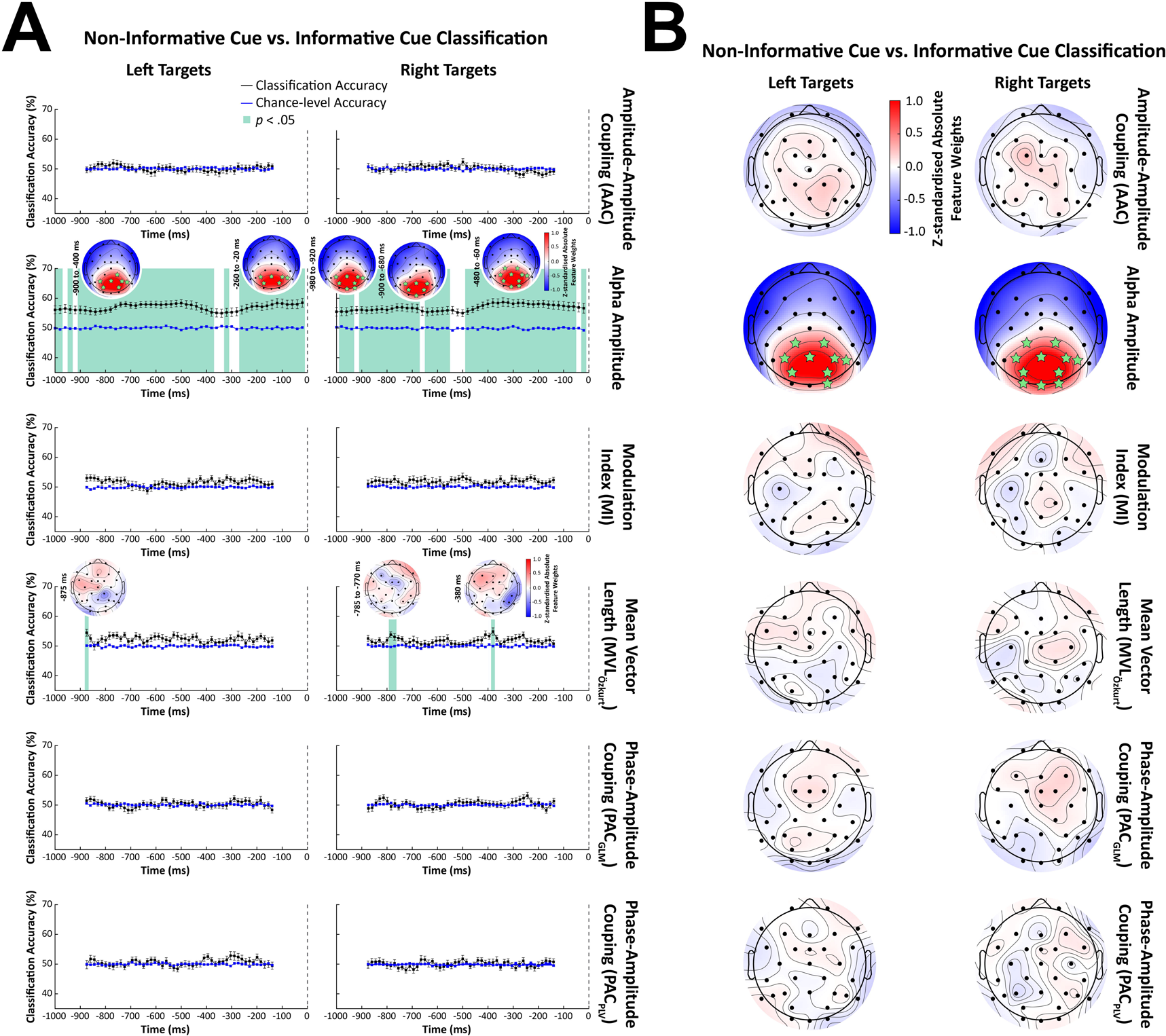
Results from spatiotemporal SVM classification and corresponding feature weight analysis based on functional connectivity metrics. **A)** Spatiotemporal classification accuracy of trials with non-informative vs informative cues across all electrodes from functional connectivity measures. The support vector machine (SVM) was trained separately on trials with left targets and trials with right targets. The black and blue lines show the actual accuracy, and the permutation test results, respectively. The green shaded region indicates the classification accuracy was significant at this time point (*p* < 0.05, corrected for multiple comparison). Error bars indicate standard error of the mean. The dashed gray line indicates target onset. Topographic plot is the z-standardized absolute feature weights at the time point indicated by the text to the left of the head. Only feature weights with a *z*-score exceeding ±0.45 are shown. **B)** Topographic maps of the *z*-standardized absolute feature weights averaged over the −1000 ms epoch prior to target onset. Only feature weights with a *z*-score exceeding ±0.3 are shown. Correlation = amplitude envelope correlations; wPLI = weighted phase lag index; PLV = phase locking value; iCOH = imaginary coherence.

### Support Vector Regression (SVR)

Support vector regression (SVR) analysis was used to determine whether the spatiotemporal pattern of EEG activity before target onset was predictive of the degree of responsive error on a given trial. As seen in Figure 5, predictions of response errors on trials with a right target and non-informative cue from spatiotemporal alpha amplitude patterns were significantly above chance at −640 ms prior to target onset. Feature weight analysis showed that the biggest contributors to prediction accuracy at −640 ms were the occipital, parietooccipital, and parietal electrodes. However, the most lateral parietal electrodes (P7 and P8) were not significant features as well as the next most lateral parietal electrode on the left (P5). The average feature weights across the entire epoch found the same electrodes to be significant contributors as well as the medial centroparietal electrodes CP1 and CP2 (Figure 5B). Note that because feature weights are only as useful as the information represented by the pattern of activity they are derived from (*e.g.,* the relative importance of electrode activity to prediction accuracy is meaningless if there is no way to predict response errors in the first place), only measures that had prediction accuracies above chance are discussed.

**Figure 5.**
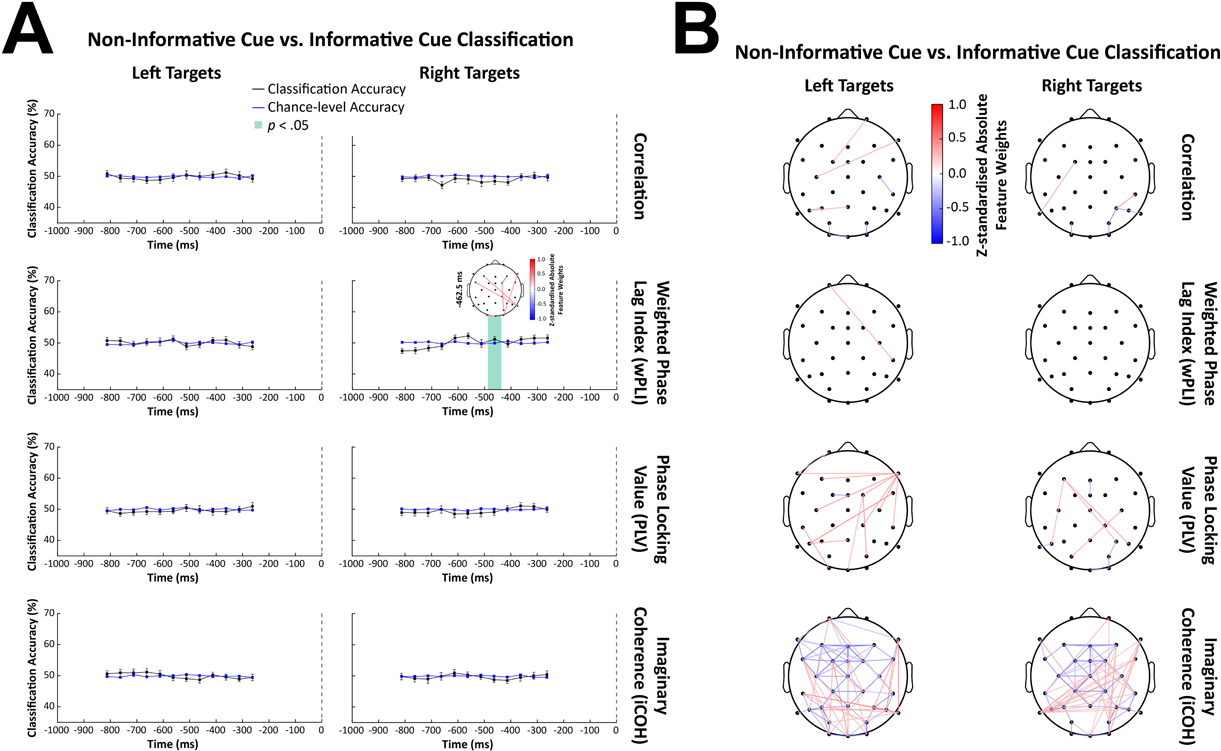
Results from spatiotemporal SVR prediction analysis and corresponding feature weight analysis based on alpha amplitude and cross-frequency coupling (CFC) measures. **A)** Spatiotemporal prediction accuracy of response errors across all electrodes for alpha amplitude and cross-frequency coupling (CFC) measures when cues were non-informative (*left*) and informative (*right*). For the non-informative cue and informative cue groups, columns are subdivided by targets on the left and targets on the right. The black and blue lines show the actual accuracy, and the permutation test results, respectively. Error bars indicate standard error of the mean. The dashed gray line indicates target onset. The green shaded regions indicate the prediction accuracy was significant at these time points (*p* < 0.05, corrected for multiple comparison). Topographic plots are the *z*-standardized absolute feature weights at or across the significant time points indicated by the text to the left of the heads. Green stars on the topographic plots are electrodes found to be significant features after correction for multiple comparisons. **B)** Topographic maps of the *z*-standardized absolute feature weights averaged over the −1000 ms epoch prior to target onset. Green stars indicate the electrodes with z-score feature weights significantly above zero after correction for multiple comparisons. Only the trial condition and measure with prediction accuracy scores significantly above chance have stars indicating the electrodes with significant feature weight *z*-scores. AAC = amplitude-amplitude coupling; MI = modulation index; MVL = mean vector length; PAC = phase-amplitude coupling.

#### Cross-Frequency Coupling (CFC)

Using the spatiotemporal pattern of amplitude-amplitude coupling (AAC) activity, predictions of response errors on trials with left targets and non-informative cues were significantly above chance at −200 ms relative to target onset (see Figure 5A). Feature weight analysis found that the biggest contributor to support vector regression (SVR) accuracy at −200 ms was the frontal electrode Fz. For the entire epoch, the most important features were the left frontocentral electrode FC1 and the frontocentral electrode FCz (Figure 5B). No other CFC metric was able to predict response errors above chance level.

#### Functional Connectivity

The only functional connectivity measure that predicted response errors above chance was the correlation metric (Figure 6A). The significant effect was only for trials with a non-informative cue and targets on the left at −562.5 ms relative to target onset. Feature weight analysis revealed that none of the electrode connections were significant contributors to the decision boundary (Figure 6B). The same was true for feature analysis over the entire epoch.

**Figure 6.**
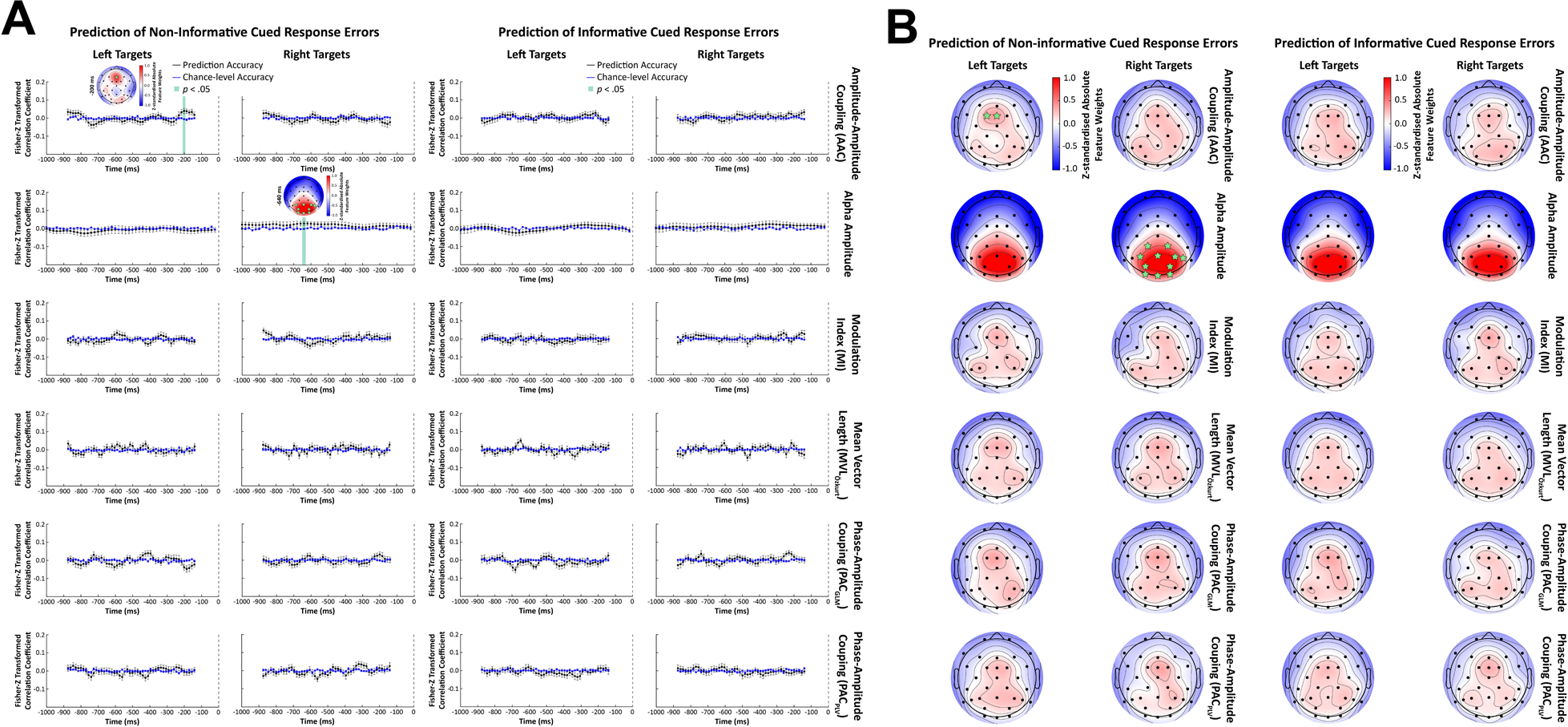
Results from spatiotemporal SVR prediction analysis and corresponding feature weight analysis based on functional connectivity measures. **A)** Spatiotemporal prediction accuracy of response errors across all electrodes for the functional connectivity measures when cues were non-informative (*left*) and informative (*right*). For the non-informative cue and informative cue groups, columns are subdivided by targets on the left and targets on the right. The black and blue lines show the actual accuracy, and the permutation test results, respectively. Error bars indicate standard error of the mean. The dashed gray line indicates target onset. The green shaded region indicates the prediction accuracy was significant at that time point (*p* < 0.05, corrected for multiple comparison). Topographic plots are the *z*-standardized absolute feature weights at the significant time point indicated by the text to the left of the heads. Only feature weights with a *z*-score exceeding ±0.45 are shown. **B)** Topographic maps of the *z*-standardized absolute feature weights averaged over the −1000 ms epoch prior to target onset. Only feature weights with a *z*-score exceeding ±0.3 are shown. Correlation = amplitude envelope correlations; wPLI = weighted phase lag index; PLV = phase locking value; iCOH = imaginary coherence.

## Discussion

In the current study, we used multivariate pattern analysis (MVPA) to test whether alpha activity, cross-frequency coupling (CFC) between alpha and gamma frequencies, or functional connectivity in alpha could better classify trials with covert spatial attention from those without. We also investigated the amount of underlying predictive information each type of alpha activity contained using MVPA. In line with the baseline sensory excitability model (BSEM) proposed by Samaha *et al* (2020), training on alpha activity alone could allow a linear support vector machine (SVM) to accurately classify trials prior to the target onset. This means that the spatiotemporal pattern of alpha activity 1000 ms before the target is presented provides the best information for discriminating trials with and without covert spatial attention. Interestingly, the same activity was only predictive of response errors in trials with a non-informative cue at a very brief time window. This too is in line with the baseline sensory excitability model (BSEM) that predicts changes in alpha amplitude affect the stimulus response and alternative responses (*i.e.,* noise) equally so that discriminability does not change, or, in current study, response errors do not improve.

We also found that the phase-amplitude coupling (PAC) metric MVL_Özkurt_ had a dissociable pattern of activity, but only for brief time windows. None of the feature weights were significant making it unclear which electrode or electrodes contributed most to determining the decision boundary although the distribution of feature weight *z*-scores in the frontal areas suggests that those electrodes contain more information than other spatial locations. Regardless, the support vector regression (SVR) analysis did not find MVL_Özkurt_ activity predictive of response errors which is counter to what we would predict about alpha-gamma phase-amplitude coupling (PAC) based on the oscillation-based probability of response (OPR) model (Zazio *et al*. 2020). In contrast, the spatial pattern of amplitude-amplitude coupling (AAC) between alpha and gamma activity was predictive of response errors on non-informative cued trials towards the end of cue-target interval which is in accordance with the oscillation-based probability of response (OPR) model (Zazio *et al*. 2020). Feature weight analysis found Fz to be a significant contributor to the determination of the regression line. For the overall time course, FCz and FC1 were significant features in the support vector regression (SVR) prediction analysis of response errors on non-informative cued trials. However, as with the alpha amplitude, amplitude-amplitude coupling activity was found to be a significant predictor of response errors only on trials with a non-informative cue at a very brief time window. Unlike alpha amplitude, amplitude-amplitude coupling (AAC) did not contain discriminating information with respect to the presence of covert spatial attention suggesting a role in the temporal anticipation for the upcoming target that provides only modest benefits to task performance.

It might seem counterintuitive that the support vector regression (SVR) results were specific to left targets or right targets when the cue was non-informative. Afterall, a non-informative cue means that participants did not know where the target will appear so they could not be influenced by the target’s location during the analyzed time window. However, the SVR analysis is using the EEG data to predict response errors which are influenced by the target’s location (see Behavioral Data section under Results). Furthermore, response errors showed an interaction between cue type and target side which might, in part, be attributed to the complex spatiotemporal pattern of alpha-related brain activity prior to target onset.

Even though we did not find as strong of evidence supporting the oscillation-based probability of response (OPR) model (Zazio *et al*. 2020), that does not mean it should be discarded. Many findings on alpha-gamma phase-amplitude coupling (PAC) are for the high gamma frequency range, something that surface EEGs are not suitable for recording due to attenuation of the electrical signals by skull and scalp. For example, Voytek (2010) found phase-amplitude coupling between theta/alpha and 80-150 Hz gamma in two participants with implanted subdural electrocorticography in visual cortical regions during visual tasks. Bonnefond and Jensen (2015) also found that the phase of alpha oscillations was coupled with the power of 80-120 Hz gamma band activity in visual sensory regions during an anticipatory pre-distractor period. Therefore, it is possible alpha-gamma phase-amplitude coupling (PAC) is important to attentional modulation, but the gamma frequency this occurs at is too fast to be recorded from outside the head.

It is important to note that while the EEG signal was filtered around the peak frequency of the aperiodic-adjusted alpha band activity, the instantaneous amplitudes extracted from the filtered signal were not aperiodic-adjusted since the FOOOF algorithm is not designed or easily adaptable for time-resolved analysis (Donoghue et al. 2020). However, we did not find the aperiodic activity to vary significantly with the experimental conditions suggesting that the current results cannot be attributed to changes in the broadband activity. At the same time, the aperiodic component was not removed from the EEG signal so it cannot be discounted entirely.

The current study only focused on activity in the alpha frequency band and its interactions with gamma band activity since it has been the major focus of research on visual spatial attention and perception. However, activity patterns within and between other frequency bands have been implicated in attentional and perceptual functioning (Clayton et al. 2015; Yuan et al. 2021). For example, previous research has found that prestimulus theta activity (4-7 Hz) is also associated with perceptual outcomes and attentional modulation (Landau et al. 2015; Fiebelkorn and Kastner 2019). In the current study, the cue-target interval was a little too short to adequately capture frequencies below alpha, but this is something that future studies can easily change by making small adjustments to the task design. Longer cue-target intervals would have the additional benefit of being long enough for time-resolved versions of other types of connectivity analysis such as those that test for directionality like Granger causality and phase transfer entropy (Cohen and van Gaal 2013; Bastos and Schoffelen 2016; O’Neill et al. 2018).

The functional connectivity metrics used in the current study are a small subset of the methods that are available. Most recent studies combining brain connectivity estimations with machine learning methods focus on various neurological conditions with much less extensive evidence on functional connectivity influencing perceptual responses (Cao et al. 2022; Sadaghiani et al. 2022). It is very possible that functional connectivity contains more decodable information about the task conditions and performance measures than these results suggest. If temporal order or coupling direction contain the most decodable information about covert spatial attention and task performance, our functional connectivity measures would not be able to capture it. Furthermore, research has found theta band activity to also be important for coordinating different attention-related brain regions (Gootjes et al. 2006; Clayton *et al*. 2015; Fiebelkorn et al. 2018; Fiebelkorn and Kastner 2019) which would explain why we found little evidence for alpha functional connectivity.

## Summary

Overall, we find more evidence supporting the baseline sensory excitability model (BSEM) proposed by Samaha *et al* (2020) than the oscillation-based probability of response (OPR) model put forth by Zazio *et al* (2020). However, the inherent limitations in the current study mean that this should not be taken as invaliding the oscillation-based probability of response (OPR) model. Instead, these results highlight the need for further investigation into the role of cross-frequency coupling (CFC) and functional connectivity in attention and visual perception. Future research can expand on our multivariate approach by investigating a wider range of frequencies whose role might have gone unnoticed due to complex spatiotemporal dynamics that are difficult to detect with traditional univariate approaches.

## Acknowledgements

This work was supported by start-up funds from the Faculty of Science and an NSERC Discovery grant (#RES0024267) awarded to Kyle Elliot Mathewson. The authors would like you thank Michael Calaresu, Jenny Le, and David Lam for assistance in data collection, and Daniel Robles and Jonathan Kuziek for analysis feedback.

## Conflict of Interest

The author declares that there is no conflict of interest that could be perceived as prejudicing the impartiality of the research reported.

## Author Contributions

KEM and SSS conceived the experiment. SSS collected and analyzed the data. KEM facilitated the results interpretation. SSS wrote the first draft of the manuscript. SSS and KEM edited the manuscript.

## Data Accessibility

The data that support the findings of this study are openly available through an Open Science Framework repository at https://osf.io/u6fgm/. Scripts to reproduce all reported analyses and figures are available through a Github repository at https://github.com/APPLabUofA/MVPA_CuedOrientTask_paper.

## Notes

https://osf.io/u6fgm/

https://github.com/APPLabUofA/MVPA_CuedOrientTask_paper

### Competing Interest Statement

The authors have declared no competing interest.

